# Activity-dependent expression of reporter proteins at dendritic spines for synaptic activity mapping and optogenetic stimulation

**DOI:** 10.1101/095984

**Authors:** Francesco Gobbo, Laura Marchetti, Claudia Alia, Stefano Luin, Antonino Cattaneo

## Abstract

Increasing evidence points to the importance of dendritic spines in the formation and allocation of memories, and alterations of spine number and physiology are associated to memory and cognitive disorders. Synaptic connections and pathways constitute the physical substrate that conveys information in the brain, and different combinations of active synaptic connections are believed to be responsible for the encoding of specific memories. In addition, modifications of the activity of such subsets of synapses are believed to be crucial for memory establishment, but a way to directly test this hypothesis, by selectively controlling the activity of potentiated spines, is currently lagging behind. Therefore it would be important to develop methods to tag active synapses for mapping functionally active connections and to selectively stimulate or interfere with active synapses. Here we introduce an approach to express light-sensitive membrane channels at synapses in an activity-dependent way by means of RNA and protein regulatory sequences. This approach is based on the local expression of reporter proteins, including optogenetic probes, at activated synapses and will allow the mapping of previously active synapses and the re-activation of the neuron only at these sites. This will allow extending the investigation of memory processes beyond the current neuron tagging technologies, whose resolution is limited at the cellular scale. Thus, it will be possible to unveil and recall the synaptic engram out of the global set of synapses.

---

Understanding the physical substrates of the mnemonic processes is one of the greatest challenges in neuroscience. Long lasting changes in the synaptic connectivity between neurons within brain circuits are generally accepted to be crucial for the establishment and maintenance of memories^1,2^. Similarities between synaptic and memory consolidation suggest shared mechanisms^3–6^, but the role of modifications acting at the synapse level still remains rather elusive^7,8^. In recent years, considerable advances in the definition of the neuronal ensembles that are activated during memory-related tasks have taken place^9,10^. On the other hand, increasing interest in defining physical connections within the brain has boosted the emergence of novel techniques to map neuron projections and synapses^11,12^. However, a way to tag and map the synaptic connections activated in response to a given stimulation, is currently missing. In fact, two aspects have so far remained orphan in the identification of synaptic engrams (i.e. set of synapses that encode a given mnemonic trace or part of it): (i) which synapses are tagged and undergo potentiation during memory encoding, (ii) what role the activity of potentiated synapses has in subsequent recalls of the encoded memory^13^.

Whereas much progress in the understanding of neural circuits has been made using light-gated channels (opsins)^9,14^, to date no direct investigation of the roles of synaptic inputs in the formation of memories has been possible using state of the art optogenetic tools. Indeed, the current spatial resolution of opsin expression is limited to whole cells but does not allow selective subcellular localization control. Cell-wide excitation does not take into account the complexity of different incoming pathways converging onto the same postsynaptic neuron^15^, and the synchronous activation of the whole cell does not represent a physiological condition. Attempts towards a more natural optogenetic stimulation have been conducted with the use of trafficking signals fused to the opsin amino-acidic sequence to enrich it at the somatodendritic or axonal level ^6,17^ but this approach still lacks sufficient selectivity. Furthermore, in none of these cases the expression of opsins was responsive to neuron activation.

Single synapse optogenetics can be achieved by two-photon or patterned illumination^18^, but this requires *a priori* knowledge of the identity of the synapses involved in the circuit in order to test their role in memory. Hence, a method to identify activated synapses is still required in order to be able to subsequently excite them. A synaptic activity reporter that could be translated locally making use of endogenous mechanisms underlying spine potentiation would be extremely useful for their identification. A special case of such new type of synaptic activity reporter would be represented by optogenetic protein probes. The local expression of opsins would allow the direct tag of activated synapses and hence their subsequent excitation by unrestricted illumination. Toward this aim, we describe here a novel tool for the expression of Channelrhodopsin variants at synapses in an input-specific, activity-dependent manner by combining rNa targeting elements and a targeting peptide tag. Their combination provides a toolkit which we term SynActive directing the local expression of reporter genes, including Channelrhodopsin. The use of SynActive will allow the visualization and manipulation of active inputs onto target neurons of interest, thereby establishing a “synaptic optogenetics” approach to recall and manipulate memory traces.

## Results

### *Arc* mRNA targeting element confers regulated translatability to an mRNA reporter

We developed a dual RNA/protein reporter to compare different RNA synaptic tags. Reporter transcripts encode a membrane-anchored fast-maturating fluorescent mCherry and bear different dendritic or axonal targeting elements (DTE and ATE, respectively); 12 copies of MS2 binding sites are inserted in the 3’UTR to visualize RNA by binding of EGFP-MS2 protein^19^. We found a minimal DTE derived from *Arc* 3’UTR^20^ to be the best candidate. *Arc* is transcribed in an activity-dependent manner and its mRNA localizes near synapses that experienced recent sustained activity; in resting conditions it is believed to be translationally repressed within ribonucleoparticle (RNP) granules^21^. *Arc* DTE determined a low level of mCherry expression in non stimulated cortical neurons, predominantly confined to the soma and the most proximal part of dendrites (Figure 1Ai), while a discrete, granule-like *Arc*/MS2 signal was detected in the soma and along dendrites (Figure 1A). In contrast to *Arc* DTE, strong or constitutive (alphaCaMKII and MAP2) DTEs drove a significant somatodendritic mCherry expression (Figure S1). KCl-induced activation of neurons expressing the *Arc* DTE construct dramatically increased mCherry fluorescence in dendrites as far as 100 μm away from the soma in as little as one hour (Figure 1Aii-ii’). mCherry fluorescence was quite uniform along dendrites, and strong somatic fluorescence was also detectable.

**Figure 1.**
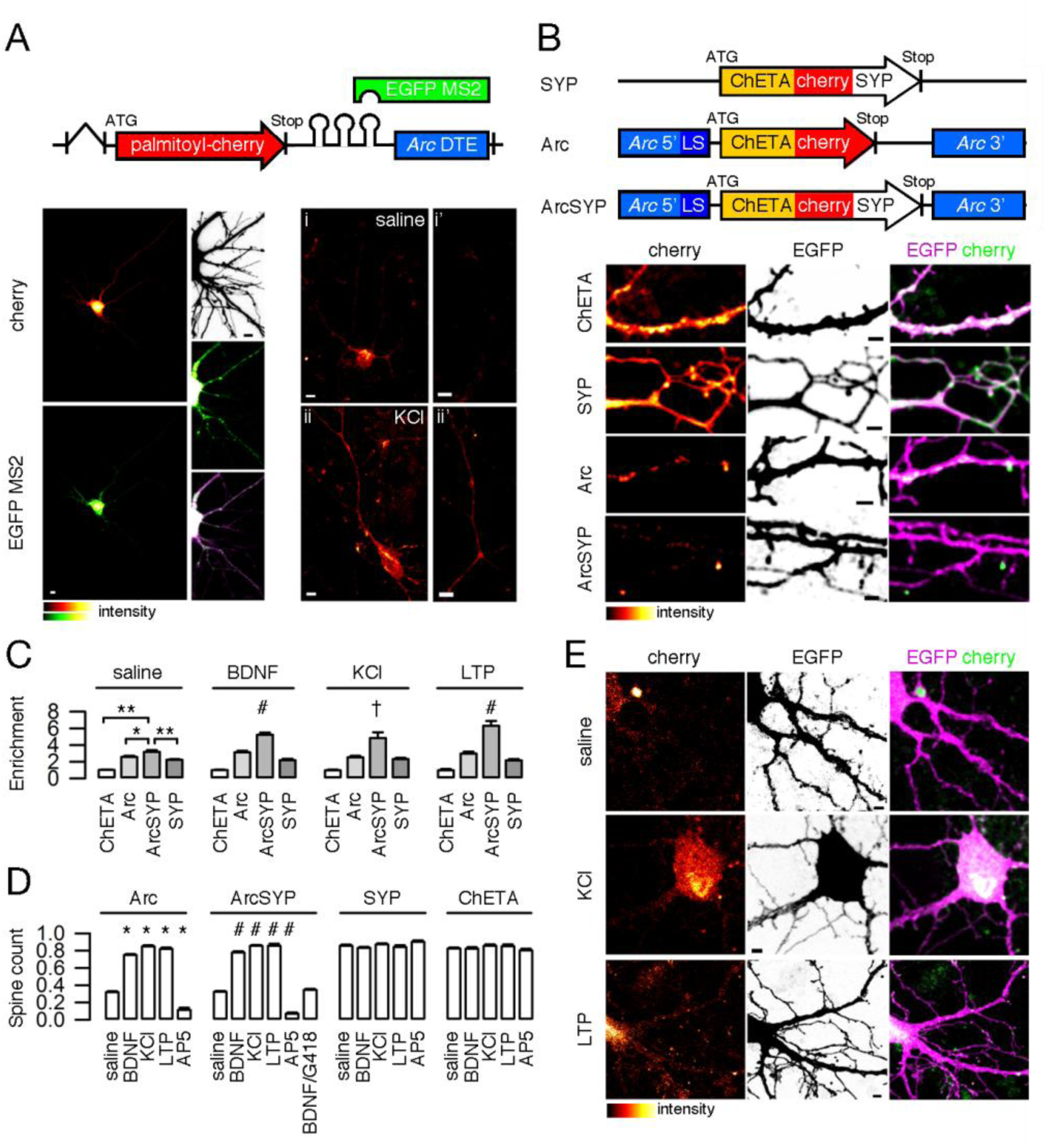
**A** Schematic construct of palmitoyl-Cherry/MS2 reporter. Left, Cherry (top) and EGFP-MS2 (bottom) distribution in living neurons under resting conditions. In the presence of *Arc* DTE, protein levels are low, and MS2/RNA signal is granular. Inset, top to bottom, neuron profile, EGFP-MS2, merge (stretched levels). Right, *Arc* DTE regulates reporter expression in response to neuron activity. Cherry expression in proximal dendrites (i-ii) and 100 μm away from the soma (i’-ii’) after 1h saline (i-i’) or 10mM KCl (ii-ii’) treatment. **B** Schematic SYP-Ch, Arc-Ch and ArcSYP-Ch constructs; throughout the figure, constructs are abbreviated as SYP, Arc and ArcSYP, respectively. Bottom, dendritic pattern of ChETA-Cherry expression (left), EGFP filler (centre) and merge (right) for unmodified ChETA-Cherry and the three constructs above. **C** Enrichment index for the three constructs and unmodified ChETA-Cherry under different stimulation conditions (see Methods). \**P*<0.01, and \*\**P*<0.001, one-way ANOVA, Bonferroni comparison of means, within group. †*P*<0.05, and #*P*<0.001 to ArcSYP-Ch, saline treated, one-way ANOVA, Bonferroni comparison of means. ND not determined. **D** Fraction of ChETA-Cherry expressing spines under different stimulation conditions, grouped for construct. \**P*<0.001 to Arc-Ch saline, and #*P*<0.001 to ArcSYP-Ch, one-way ANOVA, Bonferroni comparison of means. Differences within and between SYP-Ch and ChETA values are not significant at the 0.05 level. **E** Representative images of ArcSYP-Ch expressing neurons under different treatment conditions. Following KCl or NMDA-dependent LTP, bright ChETA-Cherry puncta are evident along dendrites. Bars are mean±s.e.m. Scale bar (A) 5 μm, (B,E) 2 μm.

### Synergistic action of RNA and protein signals for synaptic targeting of optogenetic probes

To enrich the expression of opsins at the synapses, we combined protein-and RNA-targeting sequences. We engineered fast-spiking ChETA-Cherry^22^ with the entire *Arc* 5'- and 3'- UTRs. While 3'UTR targeting elements may be sufficient for transcript localization, 5’UTR and other parts of 3’UTR generally regulate translation^23^. *Arc* 5’ leader sequence has IRES-like activity^24^, a process involved in LTP-associated synaptic translation^25^. Since ribosomes associated to spines typically lie at the junction with the dendritic shaft, we reasoned that a protein tag that interacts with postsynaptic components would improve retention and/or enrichment of the newly synthesized protein. We therefore fused to the C-terminus of ChETA-Cherry a short bipartite tag (AAAASIESDVAAAAETQV, hereafter SYP tag) composed by is the NMDAR C-terminus consensus SIESDV, and PSD95-PDZ binding consensus ETQV, which has been previously reported to enrich proteins at postsynaptic sites^26,27^.

To compare the distinct contributions of the protein and RNA instructive signals, we generated three constructs as follows: (i) *Arc* 5’-ChETA-Cherry-MS2-Arc 3’UTR (hereafter Arc-Ch); (ii) ChETA-Cherry-SYP tag-MS2 (hereafter SYP-Ch), and (iii) *Arc* 5’-ChETA-Cherry-SYP tag-MS2-*Arc* 3’UTR (hereafter ArcSYP-Ch) (see Methods for details) (Figure 1B). Neurons expressing the constructs were morphologically similar to each other or control neurons expressing EGFP alone; neither the modified SYP-ChETA nor *Arc* UTRs determined significant changes in spine number and morphology (Figure S2).

To identify the cellular compartment responsible for the translation of constructs containing *Arc* sequences, we co-expressed Arc-Ch and EGFP and compared the two signals after BDNF administration; BDNF induces a translation-dependent late form of LTP that does not directly involve neuron electrical stimulation^28^. EGFP lacked any mRNA localization element and was therefore translated in the soma. Therefore, BDNF should not directly affect EGFP dendritic levels of proteins. Following BDNF treatment, Arc-Ch signal in dendrites clearly differed from the EGFP one (Figure S3A). Conversely, SYP-Ch distribution was quite similar to the EGFP one under the same conditions. SYP-Ch fluorescence declined along dendrites in the same manner as the EGFP one, whereas Arc-Ch fluorescence was significantly higher (Figure S3B-C). This is consistent with previous observations on dendritic alpha-CaMKII: translation from reporter transcripts bearing alphaCaMKII 3’UTR was boosted by BDNF administration, increasing the protein levels along dendrites as compared to the soma^29^.

In unstimulated neurons, RNPs sequester dendritic mRNAs preventing their translation (Figure 1A and Ref^23^). Accordingly, in unstimulated cultures RNA/MS2 signal was prevalently granular along dendrites of cells expressing Arc-Ch or ArcSYP-Ch and EGFP-MS2 (Figure S4A). Following KCl treatment, the RNA/MS2 signal became much more diffuse (Figure S4A,B), indicating granule disassembling and allowing local ArcSYP-Ch translation.

We then co-expressed the three ChETA-Cherry variants with EGFP in cortical neurons to compare their sub-cellular expression pattern. SYP-Ch was evidently enriched at spines compared to unmodified ChETA-Cherry (Figure 1B); however, spines were labelled quite evenly, irrespectively of their dimension. Conversely, Arc-Ch labelled spines in a rather sparse way, with larger spines preferentially expressing ChETA-Cherry (Figure 1B). In many cases the base of the spine, rather than the whole head, was labelled most intensely, and Cherry fluorescence was also evident on the dendritic shaft (Figure 1B and 2A). ArcSYP-Ch recapitulated the sparse expression pattern typical of Arc-Ch, while more trustfully labelling spine heads (Figure 1B). Together, these observations map the essential domains that are responsible for Arc-SYP-Ch distribution: *Arc* RNA sequences determine the uneven tagging of synapses, while the SYP tag docks the protein inside the synapse.

A quantitative enrichment index (EI), the ratio of ChETA-fused Cherry intensity at the synapse to the dendritic shaft (1 to 2 μm from the spine junction), demonstrated effective ArcSYP-Ch accumulation at synapses. The EI calculated for ArcSYP-Ch was significantly higher than for Arc-Ch or SYP-Ch; all three constructs had higher EI than ChETA-Cherry (Figure 1C). ChETA-Cherry was quite uniform throughout the neuron, and, in some cases, smaller spines were not as effectively labelled as the dendrite (Figure 1B).

### Neural activity induces ArcSYP-Ch expression and selective enrichment at synapses

We next asked what regulates the expression of ArcSYP-Ch at synapses. Treatment of neurons with (i) BDNF, that mediates a late form of LTP, (ii) KCl, and (iii) NMDA, under conditions that promote spine potentiation (NMDA-induced LTP) (see Methods and Figure S5) dramatically increased the number of ArcSYP-Ch positive spines (Figure 1D-E). Conversely, NMDAR inhibition with AP5 drastically reduced the number of positive spines. Notably, the translation inhibitor G418 (geneticin)^30^ completely blocked BDNF effect on ArcSYP-Ch expression, demonstrating its dependence on novel protein synthesis. In terms of number of expressing spines, ArcSYP-Ch response to treatments was identical to Arc-Ch, whereas neither SYP-Ch nor ChETA-Cherry expression was affected by treatments that either increased or decreased neural activity (Figure 1D). Most importantly, treatments that activate neurons or induce synaptic LTP significantly increase ArcSYP-Ch EI, relative to saline treatment (Figure 1C). ChETA-Cherry and SYP-Ch were unaffected, and Arc-Ch enrichment was only modestly responsive to treatments. We ascribe this last effect to the fact that, following translation, Arc-Ch can diffuse in the membrane both onto the spine head and along the dendritic shaft; conversely, ArcSYP-Ch is retained in the spine, thanks to the SYP tag (Figure 1C).

Above-shown data demonstrate a strong dependence of ArcSYP-Ch expression on neural activity and, in particular, on synapse potentiation. In fact, LTP-inducing treatments increase the number of expressing spines, and, under control conditions, positive spines tend to be larger in dimension (Figure 1C and 2A). Spine enlargement strongly correlates with functional potentiation^31^, thus suggesting that these positive spines received a strong input stimulation from the spontaneous activity of the culture, that could amplify basal NMDAR activation^32^. Consistently, blocking NMDAR activity with AP5 drastically reduces the accumulation of Arc-Ch and ArcSYP-Ch at synapses.

**Figure 2.**
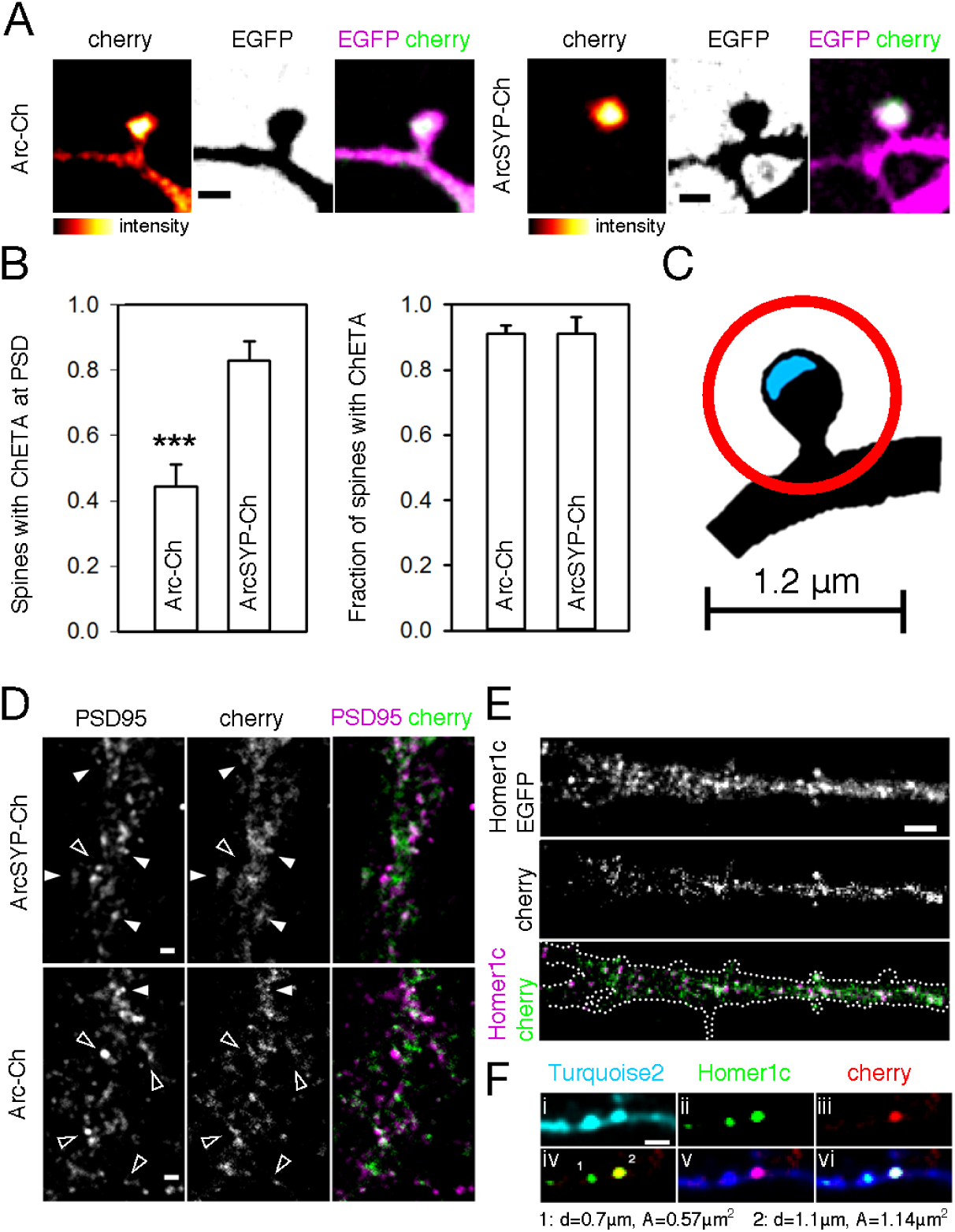
**A** Representative images of Arc-Ch and ArcSYP-Ch (cherry) expression in spines; EGFP is free in the cytoplasm. **B** Quantification of “docked”, and total positive spines following cLTP treatment (see Methods) for Arc-Ch and ArcSYP-Ch. Bars are mean±s.e.m. \**P*<0.001 two-tailed Student’s t-test. **C** Schematic drawing for the determination of “docked” versus positive but non-“docked” spines. Cherry fluorescence peaks within a circle of 1.2 μm diameter centered on the postsynaptic density (PSD - blue area) for positive spines, and on the PSD for “docked” spines. **D** Representative dendrites of neurons expressing the two constructs. White arrowheads indicate “docked” spines, empty arrowheads positive, non-“docked” spines. **E** Following a strong neuron activation (see Text), ArcSYP-Ch (cherry channel) is expressed at synapses, identified as Homer1c-EGFP puncta. Bottom, merge; dendrite profile is reconstructed with palmitoyl-Turquoise2 expression. **F** ArcSYP-Ch (cherry) is preferentially expressed at larger synapses (spine labeled as 2 versus 1 in vi) under normal culture conditions. (iv) merge of ii (green) and iii (red); (v) merge of iii (red) and i (blue); (vi) merge of i, ii and iii. Scale bar (A,D) 1 μm, (E) 5 μm (F) 2 μm.

We next asked if NMDAR-dependent LTP induction is sufficient to drive specific ArcSYP-Ch accumulation at synapses. We therefore performed double immuno-fluorescence against ChETA-bound Cherry and PSD95, one of the major components of the post-synaptic density (PSD), and compared the localization of Arc-Ch and Arc-SYP-Ch following a brief (10’) treatment of cortical neurons with NMDA under conditions that promote an increase in membrane potential and facilitate potentiation (see Methods). We considered “docked” spines those where Cherry signal coincided with PSD95, and positive, but not “docked”, spines those where Cherry intensity peaked outside the PSD, but within a circle of 0.6 μm radius centred on the PSD (Figure 2C). About half of the spines expressing Arc-Ch were “docked” (44±7%), whereas the signal of Cherry coincided with that of PSD95 in the vast majority (83±6%) of ArcSYP-Ch spines (Figure 2B-D). The total number of positive spines was the same for both constructs, and coherent with previous experiments.

We then induced a brief, strong synaptic activation of neurons by exposing neurons to NMDA under strong depolarizing conditions (60mM KCl) for 5 minutes^33^, simulating in this way high-frequency stimulation protocols used in electrophysiology that don’t make use of exogenous elevation of cAMP concentration. 90 minutes after the stimulation, strong ArcSYP-Ch expression was seen in dendrites, and the signal was neatly superimposable to that of postsynaptic Homer1c-EGFP (Figure 2E)^34^. Again, under control conditions the number of labelled spines was much smaller; ArcSYP-Ch expressing synapses had large postsynaptic densities, an indicator of functional potentiation (Figure 2F), corroborating the observation that ArcSYP-Ch positive spines, under basal conditions, were larger (Figure 1B). Experiments in hippocampal neuron cultures yield almost identical results (Figure S6): ArcSYP-Ch strongly co-localized with Homer1c-EGFP (Figure S6A), and NMDA-LTP treatment strongly increased the number of synapses expressing ArcSYP-Ch (Figure S6B). Altogether, these analyses demonstrate the activity-dependence of ArcSYP-Ch translation, as well as its preferential localization at postsynaptic sites.

### Synaptic specificity of ArcSYP-Ch expression

To demonstrate ArcSYP-Ch expression at potentiated synapses, we focally stimulated selected synapses of Arc-SYP-Ch expressing cortical neurons by means of two-photon glutamate uncaging in presence of the PKA activator forskolin^35^. To prevent potentiation of spines due to spontaneous activity under these high cAMP conditions, as well as synaptic capture, TTX was added to the medium. Focal uncaging of glutamate induced ArcSYP-Ch expression at stimulated synapses, but not at other synapses on the same dendrite or on neighbour ones (Figure 3A-C). When caged glutamate was absent, no significant change in ArcSYP-Ch intensity was observed (Figure 3B). It is unlikely that the increase in ArcSYP-Ch at the potentiated synapse is due to protein mobilization from surrounding regions, since no significant change in intensity in neighbouring spines and in the dendritic shaft was apparent. Thus, synapse potentiation is able to drive ArcSYP-Ch expression locally, without tagging non-stimulated spines.

**Figure 3.**
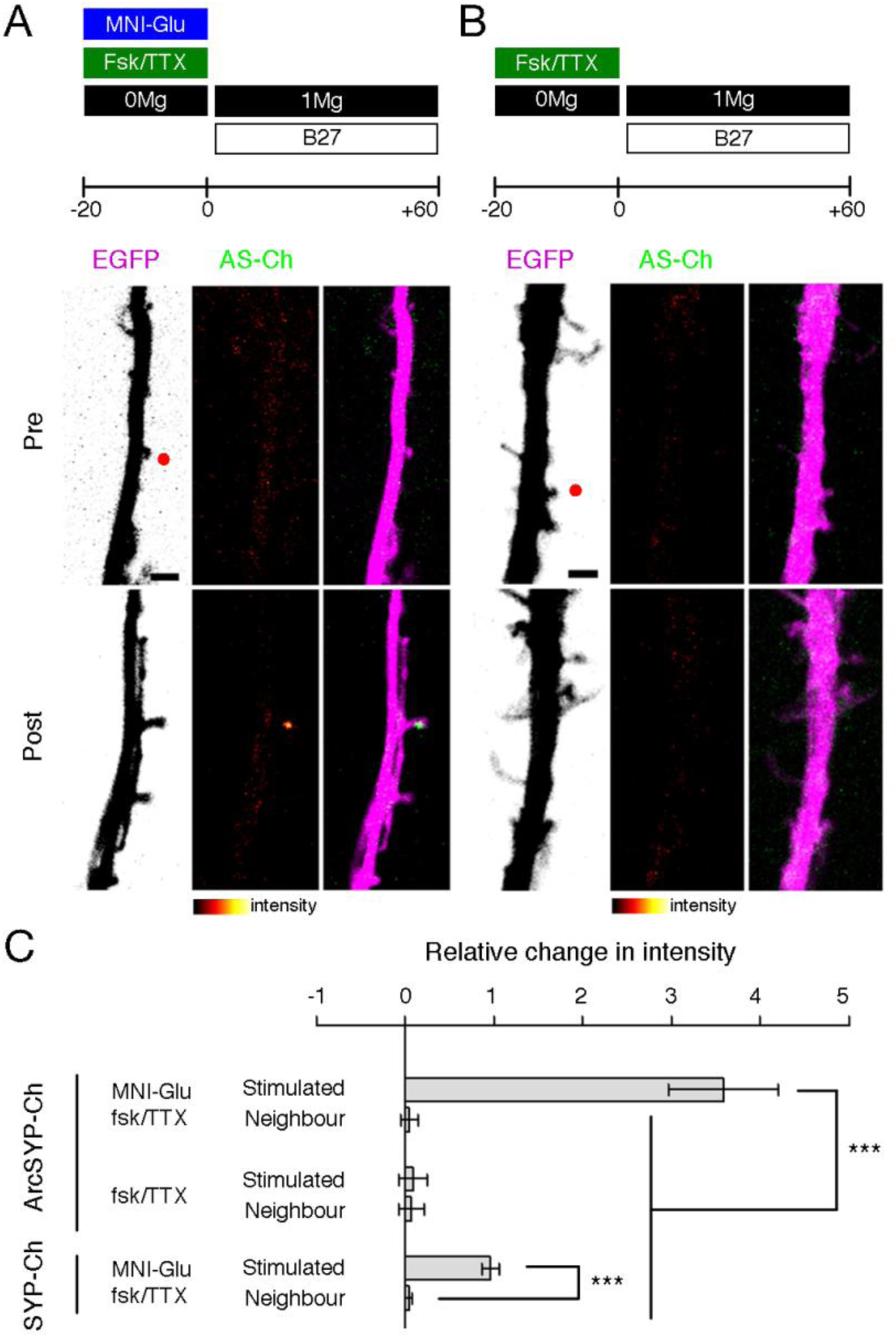
ArcSYP-Ch is specifically expressed at potentiated synapses. DIV 8-10 neurons were focally stimulated with glutamate uncaging in close proximity to selected spines. Neurons were maintained in standard Mg^2^+-free ACSF in presence of forskolin and TTX for 20’ before uncaging with (**A**) or without (**B**) MNI-caged glutamate. Following 2-photon uncaging, medium was changed to 1mM Mg^2^+ ACSF supplemented with B27. ArcSYP-Ch expression was compared before uncaging and 60’ afterwards. When MNI-glutamate was converted into its active form, ArcSYP-Ch was markedly translated in stimulated spines, but not in neighbour spines. Red dots in the EGFP channel indicate the location of 2-photon uncaging. Experimental conditions are indicated on top of images. Scale bar 2 μm. **C** Relative fold change in ArcSYP-Ch and SYP-Ch in stimulated spines and non-stimulated spines when MNI-glutamate was present or absent from batch. Relative fold change is the difference in intensity at 60’ minus the intensity at −5’, divided by the intensity value at −5’ (see Methods). Experimental conditions and construct are indicated on the left. Stimulated spine denotes the spine close to the uncaging point. \*\*\**P*<0.001, one-way ANOVA, Bonferroni comparison of means. Bars are mean±s.e.m.

Functional potentiation is paralleled by structural rearrangements that result in an observable increase in spine volume^36^, which would be reflected in an increase in fluorescence intensity as in the case of diffusible fluorescent fillers. We therefore repeated the experiment on neurons expressing SYP-Ch, which is translated exclusively at the soma level; accordingly, any change in intensity observed following LTP induction would be due to spine volume changes only. Stimulated spines underwent potentiation and increased in volume, which was paralleled by an increase in SYP-Ch signal. The observed fold changes are consistent to what previously reported^37^; however, the increase in intensity is significantly smaller than for ArcSYP-Ch (Figure 3C). We interpret these results as a further confirmation of synaptic specificity of ArcSYP-Ch expression, and they strongly imply local synthesis as the cause of the observed increase of ArcSYP-Ch at stimulated synapses.

### ArcSYP-Ch drives synaptic currents and activates neurons

In the previous sections, we established synaptic specificity of ArcSYP-Ch expression. We next asked if the locally synthesized protein is effective in driving local synaptic currents. Calcium currents are useful indicators of spine activation as a result of spine depolarization that found application both in vitro and in vivo^38–40^, and Channelrhodopsins are themselves (weakly) permeable to calcium ions^41,42^. Accordingly, a functional Channelrhodopsin is expected to cause calcium currents that could be recorded with green fluorescent genetic calcium indicator GCaMP6s^38^. We therefore restricted Illumination to imaged area by means of confocal laser scanning, and GCaMP6s fluorescence was imaged continuously.

When dendritic regions containing positive spines were illuminated, GCaMP6s fluorescence closely matched ArcSYP-Ch -bound Cherry fluorescence (Figure 4A). GCaMP6s fluorescence was much higher at the spine level than on the parental dendrite, indicating that, upon illumination, the main source for calcium influx, and therefore neuron activation, was the spine expressing ArcSYP-Ch (compare red and blue traces in Figure 4A, first column, corresponding to the temporal profiles of GCaMP6s fluorescence in the spine and in the dendrite, respectively). The temporal profile of these currents (red trace in Figure 4A, bottom panel) also closely matches a typical Channelrhodopsin current, that displays an initial peak that rapidly decays to a steady level^22,41^. This was reproducible in subsequent excitations of the same spine (Figure 4A and Figure S7D).

**Figure 4.**
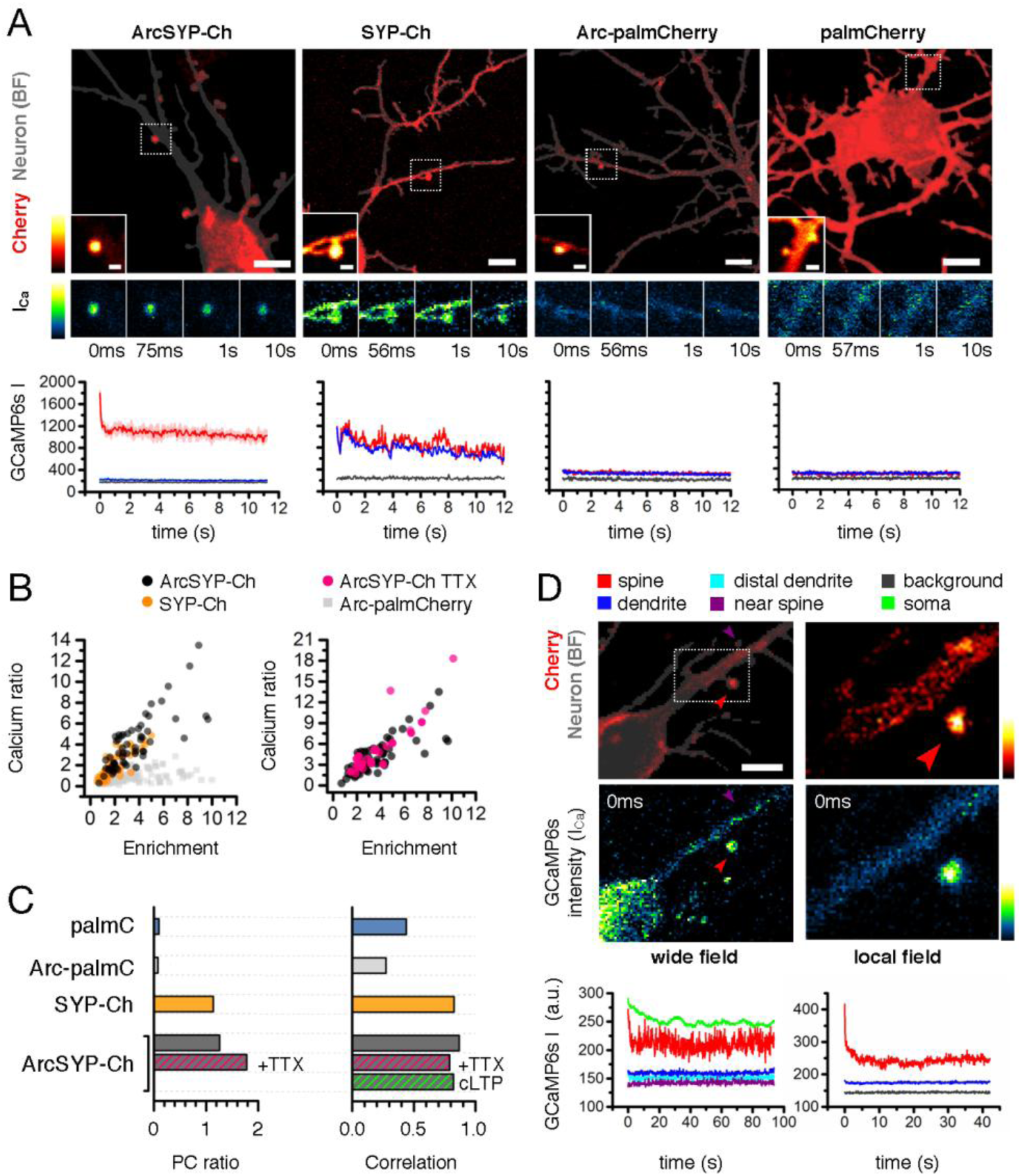
**A** Calcium currents in GCaMP6s expressing neurons cotransfected with ArcSYP-Ch, SYP-Ch, Arc-palmitoylCherrry, or palmitoylCherry (left to right). Cherry fluorescence (red) is superimposed to the neuron mask (dark grey) as drawn from the bright field channel; scale bar, 5 μm. Inset in each figure is selected spine for which calcium currents were imaged (dotted region in main figure); scale bar, 1 μm. Below are selected time points of GCaMP6s fluorescence recordings (i.e. first and second time points, and acquisitions at 1s and 10s). Bottom graphs show temporal profiles of mean GCaMP6s intensities in regions corresponding to the spine head (red trace), dendrite (blue trace) and background (grey trace). Shaded areas correspond to maximum and minimum values obtained in two consecutive repetitions. **B** Left: plot of corresponding Calcium Ratio (CR) value and Enrichment Index (EI) for analyzed spines for ArcSYP-Ch (black dots), SYP-Ch (orange dots) and Arc-palmitoylCherry (grey squares). CR is mean GCaMP6s intensity in the spine head divided by intensity in the dendrite, whereas EI is calculated as in Figure 1. Right: CR-EI plot for ArcSYP-Ch expressing spines recorded in standard recording solution (black dots) or with 3 μm TTX (magenta dots). **C** CR-EI dependence for the above constructs. PC ratio is the ratio of the coefficients of the main principal component of the distribution in the CR-EI plane (see Methods). A value close to zero indicates a very low contribution of CR to variability. Correlation is Pearson’s *r*^2^ coefficient calculated for the (EI, CR) pairs. **D** Calcium imaging of ArcSYP-Ch/GCaMP6s expressing neuron was performed illuminating in wide-field manner (left) and upon restriction of illumination to one selected spine (right, red arrowhead); scale bar, 5 μm. Below we report calcium traces for the selected expressing spine (red), a neighbouring, negative spine (purple, shown in main figure as purple arrowhead), the cell body (green trace), and distal and proximal parts of the dendrite harboring the spine (cyan and blue traces, respectively).

SYP-Ch is enriched at synapses, but it is also present along the dendritic shaft. Accordingly, calcium influx was seen synchronously both at the spine level and across the dendrites membrane (Figure 4A); notably, dendritic membrane area is considerably larger than the spines one, so even for smaller dendrites the shaft, rather that spine, is expected to be the main Channelrhodopsin electrical input onto the neuron, as it is evident from the temporal profile in Figure 4A. Neurons expressing membrane-tagged Cherry did not display comparable levels of calcium influx upon illumination; this was also true when Cherry mRNA was fused to *Arc* 5’ and 3’UTR (Arc-palmitoylCherry), in analogy to ArcSYP-Ch. The expression pattern of Arc-palmitoylCherry was very similar to ArcSYP-Ch (and Arc-Ch) (Figure 4A), but, importantly, no difference in GCaMP6s fluorescence was associated to expressing spines.

As a further confirmation, the ratio of GCaMP6s signal at the spine and at the dendrite level strongly correlated with ArcSYP-Ch EI (Figure 4B, black dots). SYP-Ch also displayed a similar dependence, but values were much smaller than for ArcSYP-Ch (Figure 4B, orange dots, and Supplementary Figure 7A). This demonstrates that the relative contribution to neuron activation strongly depends on the spatial restriction of opsin expression. Conversely, spine-to-dendrite calcium ratio was quite constant irrespectively of Cherry EI for palmitoylCherry and Arc-palmitoylCherry (grey squares in Figure 4B), and correlation values were much lower (Figure 4C).

Importantly, recorded currents for ArcSYP-Ch were not due to presynaptic activity, as both their presence and the dependence of the Calcium Ratio on EI were unaffected by TTX suppression of action potentials (Figure 4B). This implies that previously active inputs onto the postsynaptic neuron can be reactivated by the excitation of expressed ArcSYP-Ch. Furthermore, chemical LTP induction increased the number of detected spines four hours after induction, but did not modify the dependence of calcium currents on EI (Figure 4C and Figure S7B). Correlation was evident also between EI and peak value of the calcium current (Figure S7C). Calcium currents could be imaged in spines also when illumination was performed in a wide-field manner that comprised the soma, and GCaMP6s fluorescence levels indicated that even single inputs could drive significant localized inputs that were not overcome by the activation of the whole rest of the neuron (Figure 4D).

Whole-cell optogenetic manipulations that yield a synaptic response close to physiological situation have relied on presynaptic stimulation^43,44^, or coupled postsynaptic depolarization with focal glutamate release^45^. We therefore asked if the activation of Channelrhodopsin-tagged synapses would mirror a canonical activation based on neurotransmitter release. Sustained synapse stimulation activates CaMKII and determines its rapid phosphorylation that lasts for minutes^37,46^. We expressed ArcSYP-Ch or unmodified ChETA-Cherry in hippocampal neurons and determined CaMKII phosphorylation by immunofluorescence a few (7.5) minutes after optogenetic stimulation with a 450 nm optical fiber. We employed a similar pattern to theta burst stimuli used to induce LTP in the hippocampus (see Methods); to reduce background CaMKII phosphorylation, spontaneous activity was pharmacologically suppressed with TTX and glutamate receptors inhibitors for three hours prior to light stimulation (Figure 5A). During and after illumination, action potentials were inhibited with TTX. Light stimulation strongly increased phospho-CaMKII signal in ArcSYP-Ch expressing neurons compared to neurons that were maintained in the dark (Figure 5B). Light alone had no effect, as EGFP-only expressing neurons were not affected by the stimulation, and synaptic levels of phospho-CaMKII were comparable to unstimulated neurons expressing ArcSYP-Ch (Figure 5C). Importantly, CaMKII phosphorylation was specific to ArcSYP-Ch positive spines in optically stimulated neurons, as spines lacking Cherry signal had background phospho-CaMKII signal (Figure 5B,C). This parallels the physiological condition, as CaMKII activation is specific to stimulated spines^46^. Cell-wide activation of unmodified ChETA-Cherry also activated CaMKII, although the synaptic phosphorylation staining was much lower than what observed for ArcSYP-Ch (Figure 5B,C). Interestingly, phospho-CaMKII staining was also evident in the dendritic shaft of illuminated ChETA-Cherry neurons, but not in that of optically stimulated ArcSYP-Ch neurons. Thus, part of the CaMKII pool may fail to translocate from the shaft into the spine due to synchronous depolarization of spines and extrasynaptic sites; conversely, localized ArcSYP-Ch activation could induce CamKII phosphorylation and mobilization. Indeed, neurotransmitter-mediated synapse stimulation mobilizes CaMKII from the dendritic shaft and accumulates it at the spine head^47^. Accumulation is input-specific and is observed only at spines receiving direct glutamate stimulation^45,48^. We conclude that large-field optical stimulation of synaptic ArcSYP-Ch is able to simulate an input-specific excitation onto the postsynaptic neuron; conversely, whole-cell activation of ChETA-Cherry has a different outcome on the neuron response at the subcellular level.

**Figure 5.**
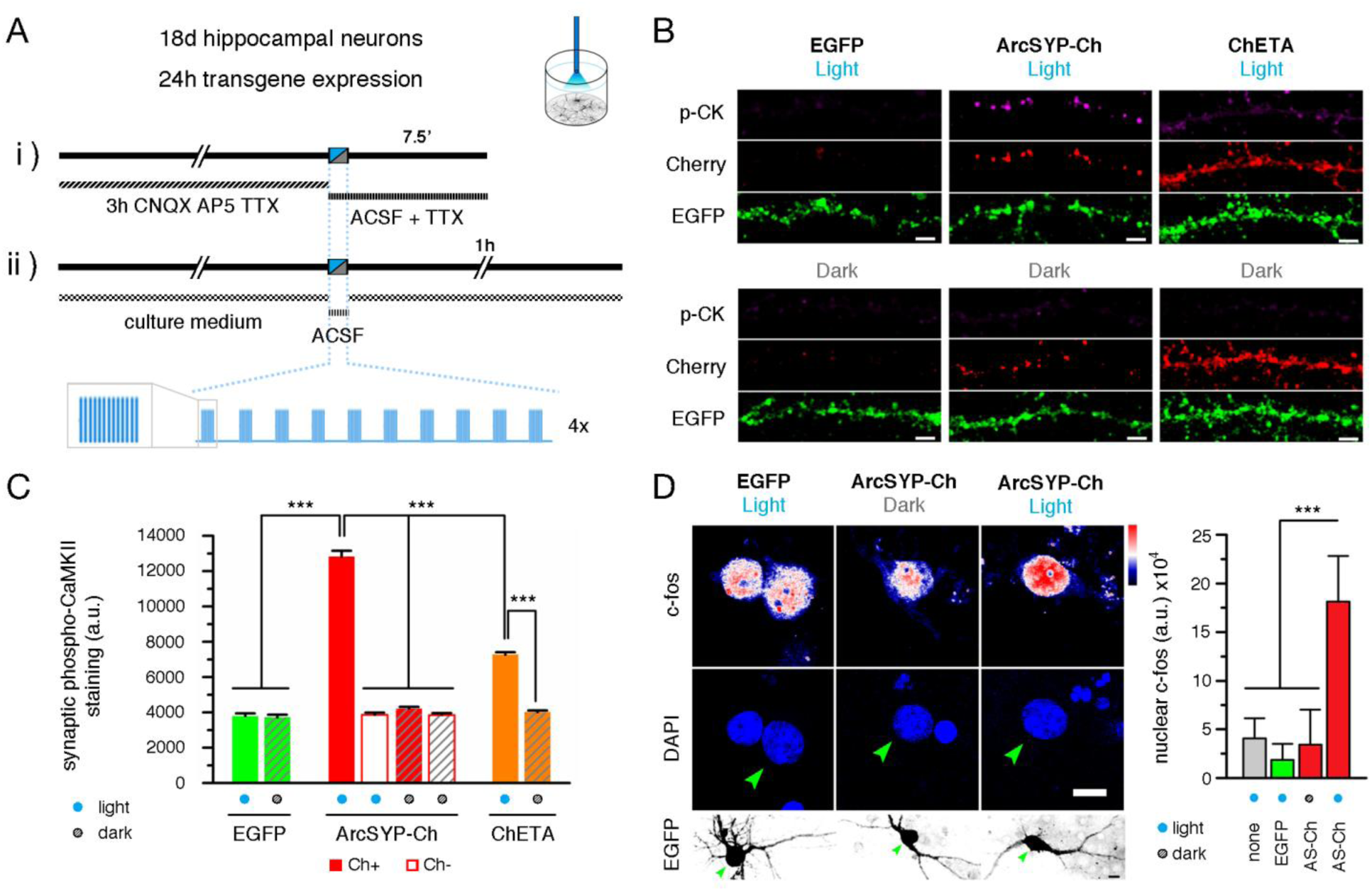
Optogenetic stimulation of ArcSYP-Ch expressing cells activates spines and induces c-fos expression **A** Outline of time course of the experiments. i) Cells were pretreated for 3 hours with CNQX, AP5 and TTX to reduce background CaMKII activation. Neurons were fixed 7.5 minutes after light stimulation and stained for phospho-CaMKII. ii) Cells were fixed and stained for c-fos 60 minutes after light stimulation. **B** Representative images of EGFP, ArcSYP-Ch/EGFP and ChETA-Cherry/EGFP expressing cells that underwent light stimulation (top panel) or were maintained in the dark (bottom panel). Panels show phospho-CaMKII immunofluorescence (p-CK), anti-Cherry immunofluorescence (Cherry) and EGFP signal. Scale bar, 5 μm. **C** Quantification of total phospho-CaMKII staining in the spine as in B. Spines in ArcSYP-Ch expressing neurons were subgrouped into Cherry positive (filled bars) and Cherry negative spines (empty bars). \*\*\**P*<0.001, one-way ANOVA, Bonferroni comparison of means. Bars are mean±s.e.m. **D** c-fos (top) and DAPI (middle row) staining of cells expressing EGFP (left) or ArcSYP-Ch and EGFP (middle and right). Cells were illuminated or maintained in the dark as indicated above. Green arrowheads indicate corresponding positions in the EGFP channel below. Scale bar, 5 μm. On the right, nuclear c-fos staining for illuminated, EGFP expressing neurons, and ArcSYP-Ch/EGFP neurons maintained in the dark is comparable to untransfected cells. Optical stimulation of ArcSYP-Ch/EGFP neurons increases c-fos expression in the nucleus. \*\*\**P*<0.001, one-way ANOVA, Bonferroni comparison of means. Bars are mean±s.d.

Last, we asked if the optical reactivation of potentiated synapses could also drive global activation of the ArcSYP-Ch expressing neurons. We therefore illuminated cultured hippocampal neurons expressing ArcSYP-Ch with 450 nm light pulses as above and we evaluated c-fos expression, an immediate early gene that is induced in neurons shortly after strong synaptic stimulation^13,49^. Stimulated neurons expressing ArcSYP-Ch and EGFP displayed evident nuclear c-fos staining one hour after optogenetic activation (Figure 5D). Conversely, c-fos staining was lower in control cells transfected with ArcSYP-Ch and EGFP that were maintained in the dark. Exposure to light alone had no effect on c-fos expression, as illuminated neurons that expressed EGFP only had lower levels of nuclear c-fos, comparable to ArcSYP-Ch transfected, not stimulated neurons (Figure 5D). Thus, optogenetic stimulation of neuronal cultures confirms that optical reactivation of synapses by large-field illumination is able to recapitulate key features of neuron-to-neuron communication.

## Discussion

The identification of active neurons has long relied on the detection of the expression of immediate early genes, which are transcribed shortly after neuron activation^49^ and form the basis for the current methods for activity mapping in the brain. Making use of this property, technologies for neuron tagging have emerged in the last decade^50^. These methods have been paralleled by investigations on neural memory circuits employing optogenetics that greatly expanded our knowledge of memory allocation^10,51^. Although highly informative, this approach is still limited spatially at the cellular scale, and therefore inevitably neglects the contribution of diverse inputs converging on the post-synaptic neuron. The current paradigm, based on theoretical and experimental data, points to synapses as the building blocks of memories Furthermore, the observation that learning different tasks involve different sets of spines further supports the idea that spines and not cells are more relevant entities for information storage in the brain^52,53^. Nevertheless, neurons receive extensive synaptic inputs from different pathways, encoding information contents as diverse as two unrelated context representations^10^. The SynActive approach presented above, that combines *Arc* regulatory sequences and protein tag, would enable to extend the investigation on memory (and other) circuits beyond the spatial scale of the neuron, allowing activity-mapping at the synaptic scale.

### Arc targeting to synapses

To refine the targeting of opsins at postsynaptic sites we combined the use of RNA targeting and regulatory sequences with a short amino-acidic tag, which guides protein localization and anchoring within the spine. Notably, we harnessed a general principle shared by endogenous synaptic proteins, i.e. anchoring and relocalization following local translation, to localize reporter proteins at spines in an activity-dependent manner. This could be most important for synaptic membrane proteins, since, although their trafficking is still largely elusive, they are likely to be synthesised in spine-associated dendritic cisternae, collectively known as spine apparatus^54,55^. Our findings are consistent with most observations implying a role for *Arc* in synapse potentiation^56,57^. Importantly, ArcSYP-Ch (as well as Arc-Ch) is preferentially found at larger spines (Figure 1B and 2F), and is expressed at focally stimulated at spines (Figure 3). Notably, we tested other mRNA regulatory sequences and DTEs, including BDNF, which is targeted to dendrites in an activity-dependent manner^58,59^. However, the BDNF splice variants that we employed (exons IIa, IIc, and VI), out of the many BDNF transcripts ^58^, were much less responsive to neural activity, either due to high basal translation (IIa and IIc) or to almost undetectable translability (VI form), consistently with recent data^60^; moreover, no evident link between expression level and spine dimension was apparent.

### Active synapse labelling

Increasing focus is centred on the functional connections and local circuits established in the brain as a result of neuronal activity^61^. Imaging techniques have been recently put forward to label synapses based on fluorescent protein complementation at the pre- and postsynaptic interface^12^, which can be implemented by making use of long-established genetic or tracing technologies to restrict the mapping of synaptic sites to determinate regions, projections or cellular types. Detecting the synapses receiving specific information is a more difficult problem to tackle; efforts in the implementation of activity sensors has made it possible to record synapse activity in response to different sensory stimulation^38,62^.

Activity alone does not imply an involvement in the storage of a defined status, as well as not all active synapses are become potentiated^62^. Recently, activity reporter SEP-GluA1, which labels synapses incorporating fluorescent AMPA receptor subunit 1 on the membrane surface, has been proposed as a marker for synapse potentiation^63,64^. Indeed, AMPA receptors are rapidly exposed on the surface of spines that underwent sustained stimulation^65,66^, which is generally accepted to be responsible for the increase in the increased currents following potentiation. However, potentiation is a complex phenomenon that comprises dissociable events, and different forms of potentiation are activated by different stimulations^67^. Some forms of potentiation do not last indefinitely, and AMPA receptors incorporation may be transient in some spines^68^; indeed, potentiation can occur without the involvement of AMPA receptors^69^. The strategy we developed can act as reporter of a late-phase, translation-dependent LTP^67^ and can be used to map potentiated synapses across a population of neurons in memory task, thus enabling researchers to identify synaptic engrams.

Unlike other strategies^70^, our approach relies minimally on the incorporation of parts of endogenous synaptic proteins; it is therefore likely to be a general method to deliver proteins of interest to active synapses, for which we propose the name SynActive strategy. A straightforward application would be to modify existing GRASP technology into SynActive-GRASP to map potentiated synapses in a given pathway. By putting the post-synaptic moiety of the GRASP methodology^12^ in the SynActive local expression vectors, one would be able to selectively express it at potentiated synapses, thus to identify them among all synapses with defined presynaptic components (e.g. given neuronal population or projecting from specific brain areas). If used in conjunction with SEP-GluA1, or other reporters of synaptic activity^68^, then, it would help understanding the dynamics as well as interplays of the different LTP phases.

### Recall of previous activity at potentiated synapses

While the importance of the integration at the cellular level is well recognized^15^, optogenetic stimulation has been limited by whole-cell expression of the opsin. In fact, cell-wide optogenetic stimulation strongly activates or inhibits target neurons; any intervention most likely results in a drastic change in action potential firing, and the only control that can be exerted is the regulation of the illumination intensity. However, physiological stimulations may not necessarily result in a change in firing rate, as graded responses deriving from subsets of synapses and local dendritic integration can also contribute to information processing. Thus, while cell-wide optogenetics can be successfully employed to drive electrical or plasticity phenomena acting on a global scale^71^, it cannot be used to control subcellular events, which are believed to be ultimately responsible for processes underlying learning and memory formation^7,72^.

Although subcellular optogenetic stimulation can be achieved by restricting the illumination pattern, this requires *a priori* knowledge of the sites to be stimulated, which are not always known. Moreover, the feasible number and sparseness of distinct illumination spots heavily depend on technological aspects. On the other hand, the biologically achieved spatial restriction of Channelrhodopsin expression presented here, would allow unbiased excitation of recently activated synapses with standard experimental setups for wide field illumination. Accordingly, when more than one spine was illuminated, significant calcium influx was only detected in spines expressing Arc-SYP-Ch, but not in adjacent ones (see Figure 4D). In fact, ArcSYP-Ch expression reflects the physiological regulation and fate of dendritic transcripts during synaptic tagging and potentiation^73,74^; thus, the use of instructive RNA sequences allows unprecedented control on the expression at synapses of Channelrhodopsin variants, and possibly of other optogenetic probes, in response to neural activity (Figure 1).

Based on our results in primary cortical and hippocampal neurons, we envisage a promising application of SynActive in the elucidation of the role of synaptic potentiation in the formation and recall of encoded memories. In fact, our experiments in neuron cultures highlight a strong dependence of spine activation on the local level of Arc-SYP-Ch expression (Figure 4). Localized calcium transients, a widely employed reporter of synaptic neuronal activity^62^, are detected upon incoming stimulation onto postsynaptic site. Our results demonstrate that the activation of ArcSYP-Ch is able to provide sufficient selectivity to generate local calcium events. In addition, single spine activation does not determine all-or-none neuron activation as cell-wide Channelrhodopsin expression would, but rather a graded response that increases in strength, as well as in complexity, as the number of expressing spines increases, even when the illumination is not restricted to a region around the spine (Figure 4A,D). Optogenetic stimulation of neurons induced synaptic CaMKII phosphorylation in ArcSYP-Ch tagged synapses, an important marker of synapse activation (Figure 5). Strong illumination of neuron cultures expressing ArcSYP-Ch also increased c-fos expression, a key marker of neuron activation that generally mirrors a sustained stimulation of synaptic inputs (Figure 5D). From the data obtained in cultures, we would expect the *in vivo* application of the presented Channelrhodopsin form to be able to induce local electrical phenomena, thus activating neurons in a weighted manner, depending on the number of tagged synapses. This would reflect their actual input drive in a determined context more physiologically, in marked contrast to what available technologies used to tag and reactivate whole neurons can achieve. Ground-breaking work in the hippocampus greatly increased our understanding of memory encoding in different areas and their role in engram formation^9,10^; however, the role of synaptic inputs remains still elusive and testing the hypothesis of a “synaptic engram”, parallel to the identified “population engram”, could benefit from the optogenetic probe presented here. This could also help unravelling some discrepancies in the literature, namely the role of CA1 in the formation and processing of memories. In fact, hippocampal CA1 cells receive multiple converging inputs whose crosstalk, following whole-cell activation in current whole-cell optogenetic protocols, is likely to result in memory occlusion^10^; moreover, parallel circuits mediate excitatory and inhibitory processing of fear recall^75^, which could complicate the analysis, when employing an excitatory tool that produces a strong, all-or-none activation that does not distinguish the location and number of incoming stimuli onto the different postsynaptic cells.

### Concluding remarks

Taking advantage of Arc RNA regulatory sequences, we were able to express a Channelrhodopsin variant at synapses undergoing potentiation, establishing a novel tool to map and reactivate these sites. While this manuscript was in preparation, a novel approach towards the development of “synaptic optogenetic” strategies was employed in a recent paper that was published ^76^; by expressing a photoactivable form of Rac1 in the motor cortex, Kasai and collaborators demonstrated that the light-induced shrinkage of recently potentiated spines severely impaired motor learning. That study emphasizes the necessity of controlling selected inputs, rather than a selected population of neurons, underscoring the interest of synaptic optogenetic approaches. However, by dramatically altering actin dynamics, such approach determined a drastic alteration of the spine structure; therefore, the interference with the memory trace could not be reverted. Accordingly, it was not possible to perform a memory recall task, as the intervention was purely destructive; thus, the sufficiency of those potentiated synaptic inputs for memory encoding remains to be addressed. To tackle the problem, one would need to re-excite those synapses, which requires the expression of a suitable protein at those synapses, in order to modify their activity. Moreover, our approach is likely to be naturally extended to any variant opsin family, thus enabling to choose to either re-excite tagged inputs or to inhibit them, in recall tasks following training. Accordingly, we show that the regulation of translation conferred by Arc UTRs is maintained when attached to a different protein coding sequence such as a membrane-tagged form of Cherry fluorescent protein (Figure 4), thereby providing a versatile tool for the identification and control over activated spines.

In conclusion, we present here a novel methodology to tag synapses in an activity dependent way, and to drive local expression of an opsin of the Channelrhodospin family. When coupled to approaches developed for the inquiry of memory circuits^9^, this will allow the bidirectional interference of the synaptic inputs involved in circuit traces and memories, a “synaptic optogenetics” approach.

## Materials and Methods

### Constructs

palmitoyl-Cherry-MS2 was generated by cloning palmitoylation sequence from GAP43 to Cherry N-terminal, whereas MS2 sequence was derived from plasmid pSL-MS2 12X (Addgene #27119)^77^. *Arc* DTE is nts 2035-2701 (NCBI NM_019361.1), in accordance to^20^. EGFP-MS2 coat protein-NLS was constructed and cloned into pcDNA3.1(+) (Invitrogen) from plasmid Cherry-MS2 coat protein-NLS (a gift from A.Marcello, ICGEB Trieste). ChETA-Cherry cDNA was PCR amplified from plasmid pAAV-CaMKII-hChR2 (E123A)-mCherry-WPRE^22^. ChETA-Cherry-SYP was generated by oligo cloning sequence GCCGCCGCTGCTTCAATTGAAAGTGACGTGGCCGCAGCTGCCGA-AACCCAGGTGTAATAA (IDT technologies) in frame to ChETA-Cherry using unique site BglII site at 3’ end of Cherry cDNA. Arc-ChETA-Cherry and ArcSYP-ChETA-Cherry constructs were generated by inserting *Arc* 5’ and 3’ UTRs before and after ChETA-Cherry and ChETA-Cherry-SYP cDNA, respectively. *Arc* UTRs were amplified from plasmid pCMV-ArcF encompassing whole 5’UTR and first 13 nucleotides of *Arc* CDS, where start ATG was mutated to ACG, and whole 3’UTR^20^. MS2 sequence was inserted downstream STOP codon before 3’UTR. Constructs were cloned into plasmid pcDNA3.1(+) (Invitrogen) under CMV promoter. EGFP was expressed from plasmid pN1-EGFP (Clontech). Homer1c-EGFP was kindly provided by D.Choquet, Institut interdisciplinaire de Neurosciences CNRS, Université Bordeaux 2. pPalmitoyl-Turquoise2 is Addgene plasmid #36209. Arc-palmitoyl-Cherry was generated from palmitoyl-Cherry cloning *Arc* 5’ and 3’UTRs in corresponding positions in analogy to Arc-ChETA-Cherry and ArcSYP-ChETA-Cherry. GCaMP6s was expressed from pGP-CMV-GCaMP6s (Addgene #40753).

### Cell culture

Primary cortex and hippocampal neurons were extracted from P0 B6126 mice as described in Ref^78^, with modifications. Following surgery and tissue isolation, tissue was triturated in cold calcium-free HBSS with 100 U/ml penicillin, 0.1 mg/ml streptomycin and _digested in 0.1%_ trypsin 100 U/ml DNase. Following trypsin inactivation in 10% FBS DMEM (Invitrogen), neurons were seeded on previously poly-D-lysine coated glass coverslips or plasma-treated poly-D-lysine coated Willco dishes. For initial plating, neurons were maintained in Neuronbasal-A medium (Invitrogen) supplemented with 4.5 g/l D-glucose, 10% FBS, 2% B27 (Invitrogen), 1% Glutamax (Invitrogen), 1 mM pyruvate, 4 μM reduced glutathione, 12.5 μM glutamate. From the following day on, neurons were grown in Neurobasal-A medium (Invitrogen) supplemented with 2% B27 (Invitrogen) 1% Glutamax (Invitrogen) 1-10 μg/ml gentamicin. Medium was refreshed every 2-4 days. For experiments in Fig.1A and S1, div 12 neurons were used. All other experiments employed div 17-19 neurons. Neurons were transfected with calcium phosphate method the day before experiment. All procedures involving animals respect Italian Ministery of Health as well as Italian National Research Council (CNR) guidelines.

### Treatments

Neurons as in Figure 1 were treated for 1h with either KCl to a final concentration of 10mM or saline added to bath. Otherwise, treatments are (i) BDNF: hBDNF (Alomone) 100ng/ml 90’; (ii) KCl: KCl 10mM 90’; (iii) LTP: 20’ in 2mM CaCl_2_/1mM MgCl_2_ ACSF followed by 10’ in 2mM CaCl_2_/Mg^2+^-free ACSF 5.4mM KCl 100 μM NMDA (Sigma-Aldrich) 20 μM glycine (Sigma-Aldrich) 0.1 μM rolipram (Sigma-Aldrich) as described in^79,80^, followed by 90’ in culture medium; (iv) AP5: 50 μM AP5 (Sigma-Aldrich) from transfection to analysis (16-20h). See also Figure S3 for temporal outline of treatments. Neurons in Figure 2E are treated with 20' 2mM CaCl_2_/1mM MgCl_2_ ACSF followed by 5’ in 2mM CaCl_2_/Mg^2^+-free ACSF 60mM KCl 100 μM NMDA (Sigma-Aldrich) 20 μM glycine (Sigma-Aldrich) (or 25’ in 2mM CaCl_2_/1mM MgCl_2_ ACSF as control) and fixed after 90’.

### Immunofluorescence

Neurons expressing Arc-Ch or ArcSYP-Ch were fixed in 2% formaldehyde 5% sucrose PBS and permeabilized in 0.1% Triton X-100. After PBS washing, samples were blocked in 1% BSA PBS, and primary antibodies anti-Cherry (GeneTex GTX59788) and anti-PSD95 (Abcam ab9909) were used in 0.5% BSA PBS. After washing, primary antibodies were detected with anti-rabbit-TRITC and anti-mouse-Alexa647 in 0.5% BSA PBS. Coverslips were mounted in Fluoroshield (Sigma-Aldrich) mounting medium. Hippocampal neurons expressing EGFP, ChETA/EGFP, SYP-Ch/EGFP or ArcSYP-Ch/EGFP for 24h were processed as above. Primary antibody was anti-MAP2 (Abcam ab5392) and it was detected with anti-chicken-Alexa647.

### Two-photon uncaging

DIV 8-10 cortical neurons were seeded on plasma-treated, poly-D-lysine coated Willco dishes and transfected the day before experiment. Neurons were maintained in Mg^2^+-free ACSF (in mM, 136 NaCl, 2.5 KCl, 2 CaCl_2_ 10 D-glucose, 10 HEPES, 2 pyruvate, 1 ascorbic acid, 0.5 myo-inositol) with 10 μM forskolin (Tocris BioSciences) 1 μM TTX (Tocris BioSciences) and, where indicated, 2.5 mM MNI-caged glutamate (Tocris BioSciences) for 20’ before uncaging. Following EGFP and Cherry acquisition, 30 pulses (720 nm, 9-13 mW at the objective lens) of 7 ms were delivered at 0.5 Hz at 0.5-1 μM from spine head. After 5’, medium was changed to 1mM MgCl_2_ ACSF supplemented with 2% B27 and the same dendrite was imaged after 60’. The mock stimulation was conducted in the same way except that MNI-glutamate was not added in the medium. Throughout the whole protocol, neurons were maintained at 37°C under humidified 5% CO_2_ atmosphere.

### Microscopy

512×512 pixels optical sections were acquired with a confocal microscope (Leica TCS SP5 on DM6000, equipped with MSD module) using an oil objective HCX PL APO CS 40.0X (NA=1.25), and pinhole was set to 1.47AU. Digital zoom was adjusted to correctly sample spines. For whole cell reconstruction z-stacks were acquired every 0.5 μm. Sequential illumination with HeNe 633, Ar 561, Ar 488, Ar 458, and diode (Picoquant, Berlin, Germany) 405 laser lines was used for Alexa647, TRITC and Cherry, EGFP, Turquoise2 and DAPI, respectively.

For two-photon uncaging, images were acquired using an Olympus FV1000 confocal module on an inverted IX81 microscope with immersion oil objective UPLSAPO 60X (NA=1.35), and pinhole was set to 180 μm. Digital zoom was set to 8x. Used laser lines were Ar 488 and HeNe 543 for EGFP and Cherry excitation, respectively. For two-photon illumination, 720 nm line was set on a tunable Chameleon Vision II Ti:Sapphire pulsed laser (Coherent, 80MHz). Green and red channels were acquired before 720 nm stimulation (-5’ time point) and 60’ after medium change (see two photon uncaging section).

### Calcium imaging

Div 7-11 cortical neurons grown on glass-bottom coverslip expressing GCaMP6s and (i) ArcSYP-ChETA, (ii) SYP-ChETA, (iii) palmitoyl-Cherry, or (iv) Arc-palmitoyl-Cherry were imaged using an Olympus FV1000 confocal module on an inverted IX81 microscope with immersion oil objective UPLSAPO 60X (NA=1.35). After red channel acquisition, selected areas were imaged with the 488 laser line at 2 μs/pixel. Depending on image size, the time step between two consecutive frames ranged from 50 to 80 ms. Neurons were maintained in standard ACSF containing 2mM CaCl_2_ 1mM MgCl_2_ 2% B27 at 37°C under humidified atmosphere. In some experiments, 3 μM TTX (Tocris BioSciences) was added to bath to prevent spontaneous firing. cLTP was induced by pretreatment of neurons in standard 1mM MgCl_2_ ACSF with 10 μM forskolin for 20’, followed by 10’ in 2mM CaCl_2_/Mg^2^+-free ACSF 100 μM NMDA 100 μM glycine 10 μM forskolin. After treatment, neurons were washed in 1mM MgCl_2_ ACSF and put back in neuron medium for 4 hours before imaging of calcium currents; during recording, neurons were maintained in standard ACSF containing 2mM CaCl_2_ 1mM MgCl_2_ 2% B27 3 μM TTX.

### Culture optogenetics

DIV 17-19 hippocampal neurons were grown on poly-D-lysine coated glass coverslips in 24wells. The day after transfection, neurons expressing ArcSYP-ChETA-Cherry and EGFP, or EGFP alone, were put in standard 1mM MgCl_2_ ACSF and illuminated with single channel PlexBright LED Module 450nm connected to an optical fiber (THORLABS, 200 μm diameter, 0.39 NA, ceramic ferrule) at 1-3 mW peak power (measured at the end of the fiber). 10 trains of 13 pulses at 100Hz were repeated at 0.5Hz; four stimulations at different positions were performed on each culture in order to evenly illuminate the whole culture area. In a first set of experiments, neurons were pre-treated for 3 hours with 40μM CNQX 100 μM AP5 1 μM TTX. Medium was changed to standard ACSF 2mM CaCl_2_ 1mM MgCl_2_ 1 μM TTX and cultures were light stimulated or maintained in the dark; 7.5 minutes after stimulation neurons were fixed for 15 minutes in 2% formaldehyde 5% sucrose PBS supplemented with 1 mM Na_2_VO_4_ 1 mM NaF to inhibit phosphatases; after permeabilization in ice-cold methanol neurons were blocked in 5% BSA 1 mM Na_2_VO_4_ 1 mM NaF PBS and subsequently incubated overnight with 1:100 mouse anti-phosphoCaMKII (Thermo Fisher MA1-047) and 1:300 rabbit anti-Cherry (GeneTex GTX59788) in 2% BSA PBS. Secondary antibodies were 1:100 anti-rabbit-TRITC, 1:100 anti-mouse-Alexa647 in 2% BSA. In another set of expertiments, neurons were light stimulated in standard ACSF 2mM CaCl_2_ 1mM MgCl_2_; after stimulation, neurons were put back into culture medium; parallel cultures did not undergo such a treatment and were maintained in the dark. After one hour, cells were fixed in 2% formaldehyde 5% sucrose PBS and permeabilized in 0.5% Triton X-100; after PBS washing, cells were blocked in 4% BSA PBS, and hybridized with 1:100 rabbit polyclonal anti c-fos (Santa Cruz sc-52) in 2% BSA 0.05% Triton X-100 PBS. Secondary antibody was anti-rabbit-Alexa647. Samples were mounted in Fluoroshield with DAPI (Sigma-Aldrich).

### Data quantification

Spine number and subclass for neurons in Figure S3 were assigned manually based on established nomenclature. Short spines with no apparent neck are classified as stubby; elongated spines whose head and neck diameters are similar are classified as thin, and spines with a defined neck and a prominent head are classified as mushroom. Filopodia were few in number across all samples and were excluded from analysis.

Enrichment index (EI) was calculated as the ratio between the Cherry average intensity on the spine region (identified using the EGFP channel) and the average intensity calculated on the dendritic shaft between 1 and 2 μm away from the spine junction, after background subtraction. For the EI calculation, only expressing spines were included in the analysis.

For two-photon stimulation experiments, spines were identified in the EGFP filler channel, Cherry fluorescence was integrated in the corresponding channel and background was subtracted. Intensity was calculated for images acquired immediately before photouncaging and after 60’ for stimulated and neighbouring spines. The relative change in Cherry intensity was calculated as the difference after and before stimulation, normalized for the initial intensity as follows: [I(+60’) − I(-5’)]/I(-5’).

Calcium Ratio is calculated as the mean GCaMP6s signal in the spine head divided by mean GCaMP6s intensity in the dendrite, once subtracted for mean background intensity. Calcium Ratio for peak current is calculated by considering just the first 3-4 recording frames before signal reached a steady value.

### Statistics

Image analysis was performed using ImageJ. Statistical analysis was performed with OriginPro v9.0. Differences between two groups were evaluated with two-tailed Student’s t-test. Residues (Figure S3) distributions were compared with Kolmogorov-Smirnov test. Multiple comparisons were made by one-way ANOVA followed by post-hoc Bonferroni test. Significance was set at α=0.05. 756 frames (21564 spines) were used for ChETA constructs expression calculation. A total of 1493 spines was analysed for EI calculation. For PSD95/ChETA-Cherry co-localization, a total of 2251 spines from 44 dendrites were analyzed. For two-photon uncaging experiments, a total of 48 samples were analysed, and a total of 118 spines were considered.

Principal component analysis in Figure 4 was performed with OriginPro v9.0. First principal component was defined as the component that explained ≥90% variance of the distribution of (EI, CR) couples. PC ratio is the ratio of the projection of the first principal component on the CR axis divided by the projection on the EI axis. Correlation is Pearson’s *r^2^* coefficient for (EI, CR) couples.

## Acknowledgement

We are truly thankful to Dr. Matteo Caleo, Istituto di Neuroscienze, Consiglio Nazionale delle Ricerche, Pisa, Italy, for making optical fiber and optogenetic setup available, and for useful discussion and suggestions. We thank Dr. S.Kindler (Uni Hamburg) for providing MAP2 and alphaCaMKII cDNA to amplify DTEs, and Dr. A.Riccio (UCL) for IMPA1 ATE cDNA. We thank Dr. A.Marcello (ICGEB Trieste) for plasmids pSL-MS2 12X and Cherry-MS2 coat protein-NLS. We wish to thank Dr. D.Choquet (CNRS Bordeaux) for making plasmid Homer1c-EGFP available. We thank C.Rizzi and N.M.Carucci (Scuola Normale Superiore, Pisa) for helping in cortex and hippocampal dissections.

